# Large field-of-view incoherent volumetric imaging in living human retina by confocal oblique scanning laser ophthalmoscopy

**DOI:** 10.1101/2021.08.05.455286

**Authors:** Wenjun Shao, Ji Yi

**Affiliations:** Department of Biomedical Engineering, Johns Hopkins University, Baltimore, Maryland, 21231, USA; Department of Ophthalmology, Johns Hopkins University, Baltimore, Maryland, 21231, USA

## Abstract

Three-dimensional (3D) volumetric imaging of the human retina is instrumental to monitor and diagnose blinding conditions. Although coherent retinal imaging is well established by optical coherence tomography, it is still a large void for incoherent volumetric imaging in the human retina. Here, we report confocal oblique scanning laser ophthalmoscopy (CoSLO), to fill that void and harness incoherent optical contrast in 3D. CoSLO uses oblique scanning laser and remote focusing to acquire depth signal in parallel, avoid the lengthy z-stacking, and image a large field of view (FOV). In addition, confocal gating is introduced by a linear sensor array to improve the contrast and resolution. For the first time, we achieved incoherent 3D human retinal imaging with >20° viewing angle within only 5 seconds. The depth resolution is ∼45 microns *in vivo*. We demonstrated label-free incoherent contrast by CoSLO, revealing unique features in the retina. CoSLO will be an important technique for clinical care of retinal conditions and fundamental vision science, by offering unique volumetric incoherent contrasts.

## 1. Introduction

Imaging the human retina is unique in its ability to create 3D microscopic images through the clear ocular media, and is significant for its translational values. While the imaging mechanisms differ among various techniques, 3D volumetric imaging in the living human retina has almost exclusively relied on coherent scattering by optical coherence tomography (OCT), leaving incoherent contrasts undetectable. On the other hand, incoherent optical contrast offers a range of valuable structural and functional information. For example, fluorescein angiography (FA) uses incoherent exogeneous fluorescence to detect retinal vascular leakage that has been and continues to be a gold standard in diagnosing retinal vasculopathy [1]. Multiple scattering and differential phase contrast enhance the contrast for transparent cells [2–5]. Autofluorescence underlies molecular retinal imaging [6,7].

Despite the above important use of incoherent contrasts, we are lacking tools to image incoherent contrasts in 3D, and currently present them in two-dimensional (2D) fashion based on fundus photography or the point scanning scheme. The prevalent fundus photography can give easy access for imaging both incoherent scattering and fluorescence [1,8,9]. However, it lacks the ability of three-dimensional imaging. Scanning laser ophthalmoscopy (SLO) and confocal SLO (cSLO) use a scanning laser [10,11] to improve the resolution and contrast. However, they underfill the pupil, equivalently limit the numerical aperture (NA), and the axial resolution is in the order of several hundred microns [12,13]. Adaptive optics SLO (AOSLO) pushes the resolution limit by fully utilizing the pupil size and by correcting wavefront aberration. Diffraction-limited resolution can be achieved in the order of 1 micron in the lateral plane and 30-40 microns in depth [14]. Yet, the point scanning approach requires extensive Z-stacking to compile a volumetric dataset [15–20,14], which is time-consuming and challenging in practice. In addition, the field of view (FOV) in AOSLO is typically limited to only 1° to 2° viewing angle.

To enable 3D incoherent retinal imaging in the human eye, we report a confocal oblique SLO (CoSLO) to collect depth signals in parallel so that z-staking is completely avoided. We essentially reengineered the concept of single objective light sheet microscopy (SOLSM) [21,22] with human ocular optics, where the ocular lens serves as a low NA objective lens. There are two major innovations that enable our work. First, a demagnifying optical design was implemented to overcome the limitation of small NA for the dilated human eye, and still achieve tomographic imaging with practical light collection efficiency [23] [24]. Second, a confocal detection was implemented to reject the diffusive light and improve the image contrast. We descanned the oblique laser line onto a linear detector array, so that the A-line depth signal is acquired simultaneously. Each photon detector served as a confocal gating to reject diffusive light.

For the first time, we demonstrated incoherent volumetric imaging over a large FOV of 6 × 6 mm^2^ area (*i*.*e*. ∼20° viewing angle) in the living human retina. Without the need for Z-stacking, the entire volumetric data can be acquired within 5 seconds. The axial resolution is ∼44.5 µm, 4-times better than that of SLO and approaching to AOSLO [12,13]. Different FOV centered at macular region and optic disk were demonstrated. We also showed that spectral contrast can be achieved with minimum change of the optics. With the capability of imaging incoherent contrasts in 3D, CoSLO is a significant development to reveal unique retinal features in the human retina.

## 2. Methods

### 2.1 Principle of CoSLO incoherent volumetric imaging

The concept of the CoSLO is illustrated in Fig. 1(a). Human ocular optics is used for both the oblique illumination and detection. By scanning the oblique laser line (A-lines) along either the fast and slow axis, an B-scan is created in the retina. The incoherent A-line signals are descanned by the same scanners such that they can be mapped onto a stationary line, also in an angle with respect to the optical axis. A tilted remote focusing system then images the descanned stationary line image with a linear detector to record an A-line. Because the depth-resolved incoherent A-line is acquired in parallel, no Z-stacking is needed as compared to the point scanning scheme. Therefore, the data throughput is dramatically improved, facilitating incoherent volumetric retinal imaging.

**Fig. 1.**
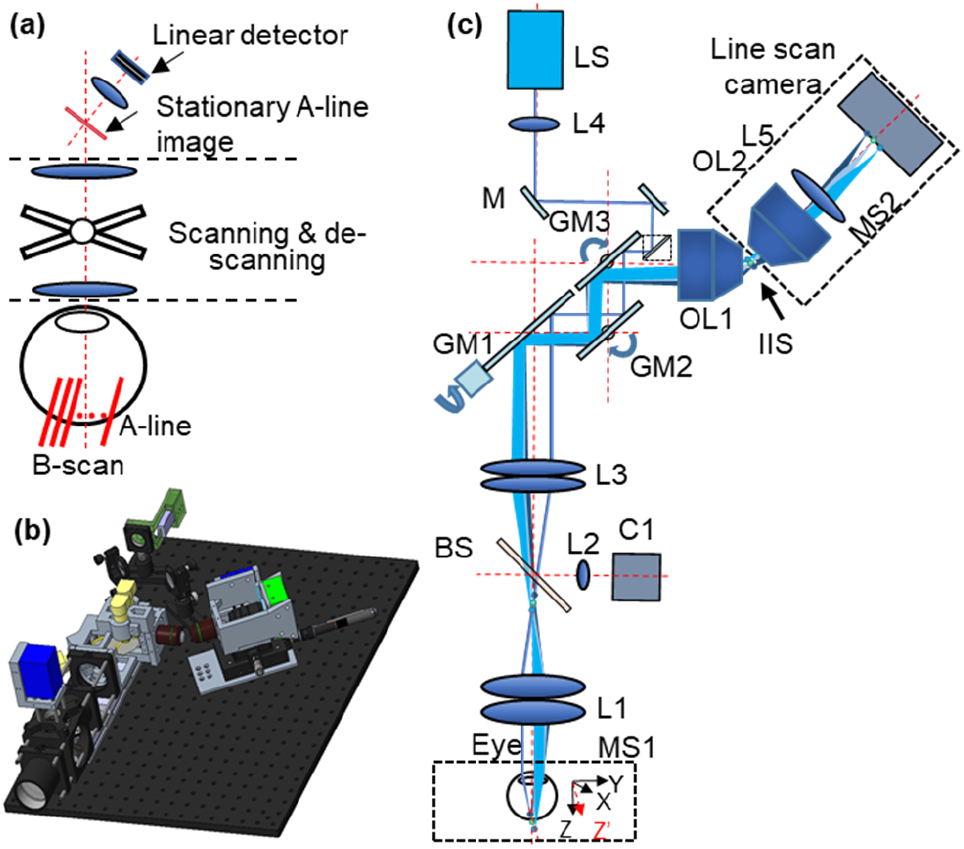
The schematic and 3D model of the CoSLO system. (a) The concept of the scanned light sheet in the eye by using CoSLO; (b) The 3D model of the CoSLO setup. (c) The system schematic of CoSLO: LS, light source; L: lens; M: mirror; GM: galvanometer mirror; MS: motorized stage; IIS: intermediate image space; OL: objective lens; BS: beam splitter.

### 2.2 System description

The 3D model and the schematic of the proposed CoSLO setup are shown in Fig. 1 (b) and c). Two light sources (LS) can be switched between near-infrared (NIR, SLD830S-A20W, Thorlabs) and visible light (SuperK Extreme OCT, NKT photonics) with other components unchanged. The NIR channel has a center wavelength of 830 nm with 55 nm bandwidth, and the visible channel has a center wavelength of 600 nm with 35 nm bandwidth. Both of them has micron-level coherent length to be approximated as incoherent imaging.

After the light was collimated by (L4: f = 10 mm), a pair of mirrors (M1&M2: PF10-03-P01, Thorlabs) was used to adjust the attitude and orientation of the beam. A right-angle mirror (M3: MRA05-F01, Thorlabs) was moved by a translational stage so that the offset of the beam with respect to the optical axis could be adjusted. To reduce the number of relay lens and the aberrations, a virtually conjugated galvanometer pair (VCGP) consisting of three galvanometer mirrors (GM1& GM2: QS-20X-AG, Thorlabs; GM3: QS-20Y-AG, Thorlabs) was used as the slow-axis mirror and the fast-axis mirror [25]. The pupil of the slow-axis mirror (GM2, GM3) was coincident with the fast-axis mirror (GM1). The exit pupil of the VCGP was then mapped onto the pupil of the human eye by a relay lens group (L3: Two ACT508-200-A, Thorlabs; L1: Two ACT508-100-A, Thorlabs). The scanning protocol is shown in [Supplement 1, Fig. S1]. As the dilated human eye usually has a diameter of ∼7 mm, the offset of the illumination is adjusted to ∼3 mm away from the center of the pupil to maximize the oblique angle of the excitation line in the retina. The beam diameter was ∼0.5 mm when projected onto the cornea of the eye. The approximated focal length for the human eye is ∼16 mm according to its optical power of 62.3 D [24]. Given the offset of the beam and the focal length of the eye, the oblique angle of the excitation line created on the retina is estimated to be ∼10°. For the NIR light, the beam waist and the Rayleigh range of the oblique Gaussian beam that focused on the retina were estimated to be ∼ 8.5 µm and ∼270 µm, respectively. As for the visible light, the beam waist and Rayleigh range are approximately 6 µm and 196 µm, respectively. The beam splitter (BS: BP208, Thorlabs) and lens (L2) were used for a pupil camera (C1) so that the position of the eye could be adjusted accordingly by a motorized chin rest (MS1). The photograph of the device construct is in supplemental Fig. S2.

The scattered light from the retina was directed through the same relay lens group into the VCGP unit, where the oblique scanning laser was descanned to be stationary in the intermediate imaging space (IIS).

One challenge with CoSLO is that human ocular optics has low NA ∼0.2, in contrast to high NA objective lens in SOLSM [21,22]. The intersection angle between excitation and detection is small. If the image of the oblique laser were to maintain the same 10° excitation angle with unity magnification, the remote focusing system would have dismal collection efficiency [27].

To overcome this challenge, we used a demagnifying design from retina to IIS, so that the angle of the stationary line image can be increased to be ∼32° with M<1 [28] by

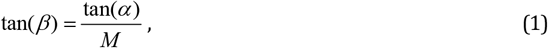

where β is the angle of the stationary line image with respect to the optical axis of OL1, α (10°) is the angle of the oblique excitation laser line on retina, and *M* is the lateral magnification from retina to IIS. *M* is designed to be 0.28 by the human ocular optics, L1, L2, and OL1 (UplanFL N 20 × /0.75, Olympus), and β is ∼32°.

Another innovative design in CoSLO is to use confocal detection scheme in the remote focusing system by utilizing a line sensor array, instead of planer cameras. As the line width of the linear photon detector was narrow, it served as the slit aperture and effectively reject out-of-focus signal [supplemental Fig. S3]. The tilted final imaging system consisted of the objective lens (OL2, LMPL50XIR, Olympus), camera lens (L5: MVL50M23, Navitar), line scan camera (OCTOPLUS-PF-SYST, Teledyne e2v), and the motorized stage. As the excitation line has an angle with respect to the Z-axis, the volume data can be transfer to a standard cartesian coordinate by an affine transformation [27].

### 2.3 Human retinal imaging procedure

The institutional review board has reviewed and approved the study. The study was Health Insurance Portability and Accountability Act-compliant and adhered to the tenets of the Declaration of Helsinki.

One drop of 1% Tropicamide was used to dilate the pupil. As the proposed confocal detection scheme allows the detection of individual A-line at any scanning position, we designed a cross scanning pattern in the alignment mode to display both the X-Z and Y-Z simultaneously. After focus adjustment and beam alignment at the pupil, satisfactory image quality can be obtained in both cross-sectional images. Then a raster scanning is used to acquire the volumetric data.

The powers on the cornea for the NIR and visible light channels were 200 µW and 60 µW, respectively. The power was well within the maximum permissible exposure power that allowed for laser safety [29]. The line acquisition rate was set to 50K Hz, and the fast axis has a duty cycle of 0.8.

## 3. System resolution

### 3.1 Theoretical framework for CoSLO

The theoretical expression for CoSLO resolution can adopt the framework for confocal laser scanning imaging with a generalization in 3D. The illumination intensity *I*_*i*_ at the sample can be written by [30]

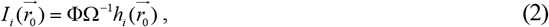

where Ф is the source power intensity; *h*_*i*_ is the illumination point spread function; and 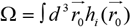 is the normalization factor. 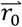 is a spatial vector in the sample space. The signal generated from the sample under *I*_*i*_ is

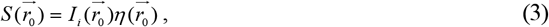

where 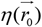 is the local coefficient in the unit of *per length*, determining the incoherent signal strength. The image formation at the intermediate imaging space between OL1 and OL2 is given by

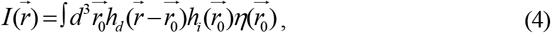

where *h*_*d*_ is the detection point spread function (PSF).

The remote focusing system with a linear detector effectively creates a virtual confocal array along the illumination laser line, where each sensor unit (*e.g*. each pixel in a line scan camera as used here) serves as a confocal aperture to gate and integrate the signal. At each pixel, the signal Γ is given by

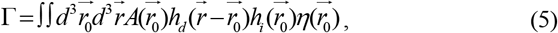

where 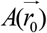 is a unitless aperture function representing the pixel area in 3D space. Because the remote focusing system is in an angle so as the virtual confocal array, 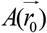 is a 3D function. Now we consider that the scanning and de-scanning mechanism equivalently translates the sample by 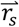. The image formation is then

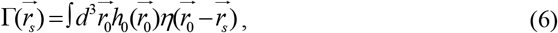

where we re-define our system PSF

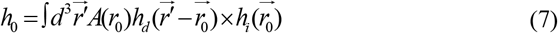

An examination of *h* finds that 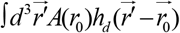 is the convolution of the virtual confocal aperture function *A* with the detection PSF *h*_*d*_, equivalently imaging the sensor array to the sample space. Equation 7 will be used for the following numerical simulation to characterize theoretical resolution.

### 3.2 Numerical simulation of the system resolution

A Fourier domain model is established to calculate the theoretical resolution as shown in Fig. 2(a). The coordinate is defined by a wave vector 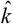,

**Fig. 2.**
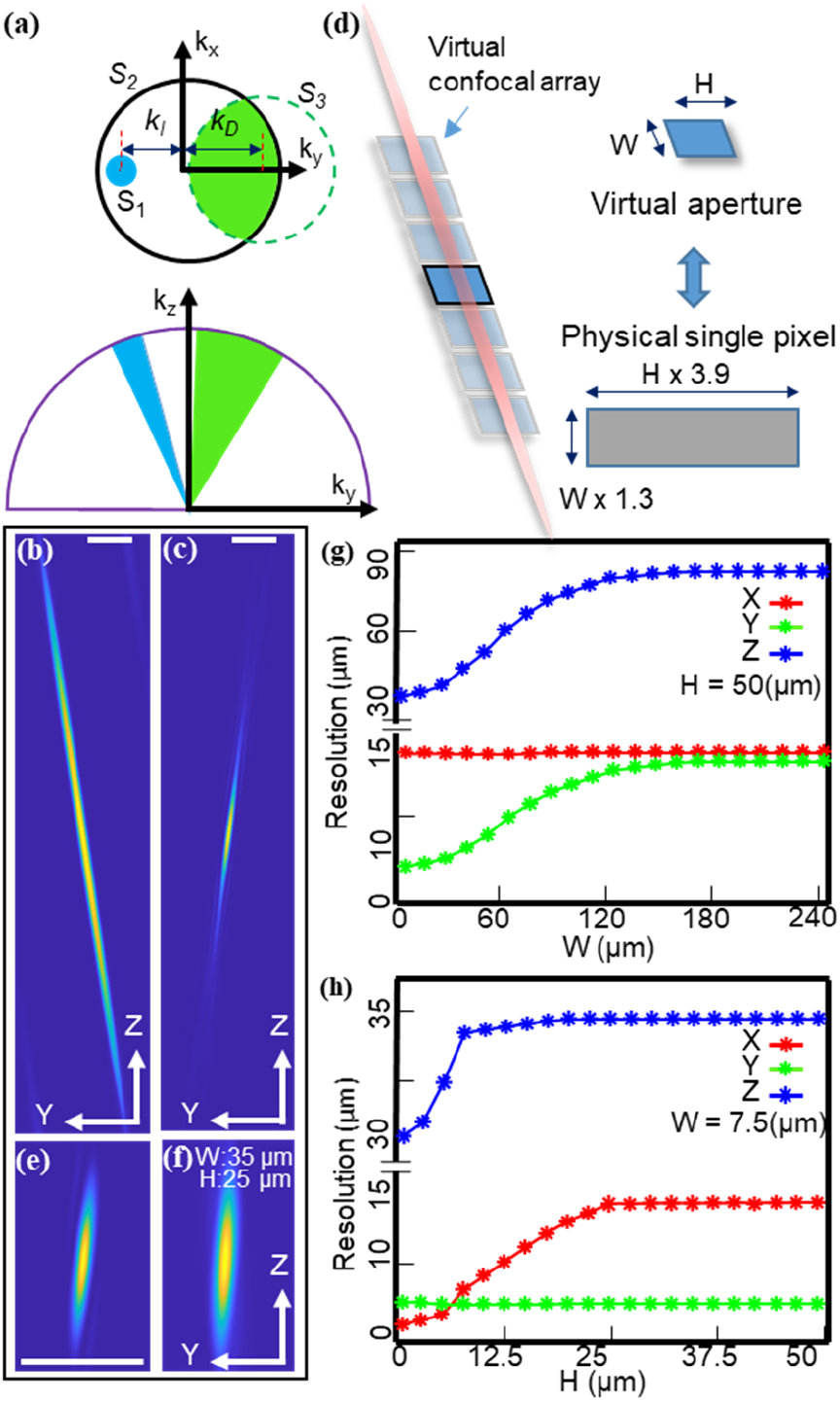
The Fourier domain model of the CoSLO. (a) The 3D frequency support of the illumination and detection in the 3D Fourier domain. The blue area S_1_ represents the illumination. The black circle S_2_ represents the spatial frequency range by the dilated human eye. The dashed green circle S_3_ shows the mapping of OL2 detection range. The offset is a result of the angled alignment for the remote focusing system. The intersection area of S_2_ and S_3_ represents the spatial frequency range of the detection. (b-c) The Y-Z cross-sections of the illumination and detection PSFs. (d) The illustration of the virtual confocal array aligned with the oblique illumination path. The virtual aperture is a demagnified image of actual sensor pixel. (e-f) The Y-Z cross-sections of the system PSFs with infinite small and finite aperture size. (g-h) The change of the resolution under varying height, *H*, and width, *W*, of the aperture. The scale bar is 100 µm.

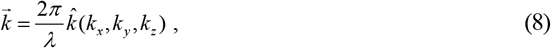

where 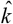 is the directional vector with unit magnitude, *λ* is the wavelength. For simplicity, the center wavelengths of the NIR and visible light in the two channels were used for calculation.

We now can describe the 3D frequency support for CoSLO illustrated in Fig. 2 (a), and specify the spatial frequency ranges of the illumination and detection. The blue circle *S*_*l*_ and represents the spatial frequency range for the illumination, and *k*_*I*_ represents the offset of the oblique illumination. Thus, the special frequency range for the illumination is identified as:

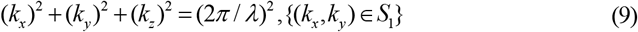

Similarly, the black circle *S*_*2*_ represents the spatial frequency range allowed by the dilated human eye (NA=0.2). The dash green circle *S*_*3*_ shows the equivalent spatial frequency range of the OL2 that is mapped on the pupil, and *k*_*D*_ represents the tilted remote focusing system. Because of the angled alignment between OL1 and OL2, only partial frequency range at the back pupil of OL1 can be collected. The overlapping area of the *S*_*2*_ and *S*_*3*_ describes the detection range, which is given as:

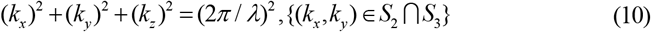

The specific calculations of the spatial frequency ranges and the frequency offsets can be found in [Supplementary 1].

After specifying the 3D frequency support, a 3D Fourier transformation was performed to obtain amplitude spread function (ASF). Then the point spread functions (PSFs) are the squared magnitude of ASFs. The PSFs of the illumination and detection (*h*_*d*_ and *h*_*i*_) are shown in Fig. 2(b-c), respectively.

Next, we consider the confocal gating in CoSLO. According to Eq. (7), the convolution between *A* and *h*_*d*_ equivalently projects a confocal array to the sample space. Because the remote focusing system is aligned to refocus the oblique line image onto the linear array, the virtual confocal array is then conjugated with the illumination beam, as illustrated in Fig. 2(d). Each virtual confocal aperture in the array is considered independent, and defined by 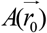. We used *W* and *H* to quantitate the geometry. Note that *W* and *L* are the size of virtual aperture which can be scaled to the physical size of actual sensor pixel by the system magnification (Supplement 1).

According to Eq. (7), when *A* is infinitely small approaching a Dirac delta function, the system PSF (*h*_*0*_) is a multiplication of *h*_*i*_ and *h*_*d*_ resulting in the diffraction-limited resolution, shown in Fig. 2(e). Larger confocal aperture size, either *W* or *H* will have relaxed resolution as expected (Fig. 2(f)). Figure 2(g) describes the change of system resolution with varying *W* at the fixed *H* = 50 µm. The axial resolution deteriorates when the *W* increases along the excitation light path. At the same time, Y resolution also becomes worse, since *W* will project to Y axis. Figure. 2(h) shows the change of the system resolution with varying *H* at *W*=7.5 µm. The axial resolution is maintained within 32-35 µm, and increasing *H* only impacts X resolution. In our setup, the line camera pixel size (200 x 10 µm) is equivalent to *H* = 51 µm, and *W* = 7.7 µm. Then the theoretical resolution is 13.5, 5, and 35 µm in X, Y, and Z.

## 4. Experimental results

To characterize the resolution of the proposed CoSLO, a schematic eye described in our previous publication was imaged [28]. A layer of agarose gel with an average thickness of ∼680 um was adhered inside the schematic eye to mimic the human retina. By imaging the microspheres with a diameter of 3.1 um that was immobilized in the gel, the resolution, as well as the field of view (FOV), can be characterized. The actual depth dimension of the agarose gel was confirmed by OCT [31] [Supplement 1, Fig. S5]. Figure 2(a-c) are three cross-sectional projections of the incoherent volumetric data on the agarose gel with NIR illumination. (Please see the flythrough of the volume data in Visualization 1). The circular boundary in Fig. 2(b) indicates the 3D-printed mold to confine the gel thickness. The diameter of the mold is 10 mm, demonstrating the large FOV by our CoSLO. Vignetting starts to appear at the edge of the FOV, limiting more in the Y direction than X. The bright signal from the bottom of the gel is the inner boundary of the schematic eye model. The overall optical magnification was calibrated to be ∼1.2 and ∼3.5 for the depth and lateral directions, respectively. The zoom-in view of the squared area in Fig 2(b) is shown in Fig. 2(d). Figure 2(e) is the X-Z cross-section taken from the area between the dashed lines in Fig. 2(d). The bright bottom in Fig 2(e) is the inner boundary of the schematic eye model. As the thickness of the agarose gel is with in twice the Rayleigh range, Fig 2(e) exhibits constant image quality in the depth direction. The profiles of three representative microspheres are shown in Fig. 2(f). To quantitatively analyze the resolution over the full FOV, the full width at half maximum (FWHM) along each direction of different microspheres was calculated. As shown in Fig 2(g), the resolutions along X, Y, and Z direction are 11.3±1.1 µm, 15.4±1.3 µm, and 44.5±3 µm, respectively.

We also evaluated the effect of confocal gating in CoSLO, and demonstrated that shorter sensor heights (*H*) effectively suppress diffusive background and improve image contrast [Supplement 1, Fig. S3].

With the resolution characterization, we proceed to image the living human retina. Two health human volunteers participated in this study. To demonstrate the FOV of the CoSLO, volumetric data covering an area of ∼6 × 6 mm^2^ in the retina, i.e., 20° viewing angle, was first acquired using the NIR channel (Fig. 3a-3c). The large FOV enables imaging across the optic nerve head (ONH) and macular region. The entire volumetric data has a pixel density of 512 × 512, acquired in ∼5s by a 50kHz A-line rate. Depth-resolved signal was acquired in parallel, as compared to sequential z-stacking in conventional point scanning methods. Figure 4(a) and 4(b) are maximum intensity projections (MIPs) which were taken from around 0-50 µm, and 120-170 µm from the retinal boundary that indicated by the yellow line in Fig. 4(c). Thanks to the depth sectioning ability of CoSLO, different features can be observed. In Fig. 4(a), nerve fiber bundles in the inner retina closed to ONH, retinal vasculatures are shown; while Fig. 4(b) shows the shadows of the vasculatures, as well as spotty signals in the background. The cross-sectional view in Fig. 4(c) also demonstrated distinct features along the depth of the retina.

**Fig. 3.**
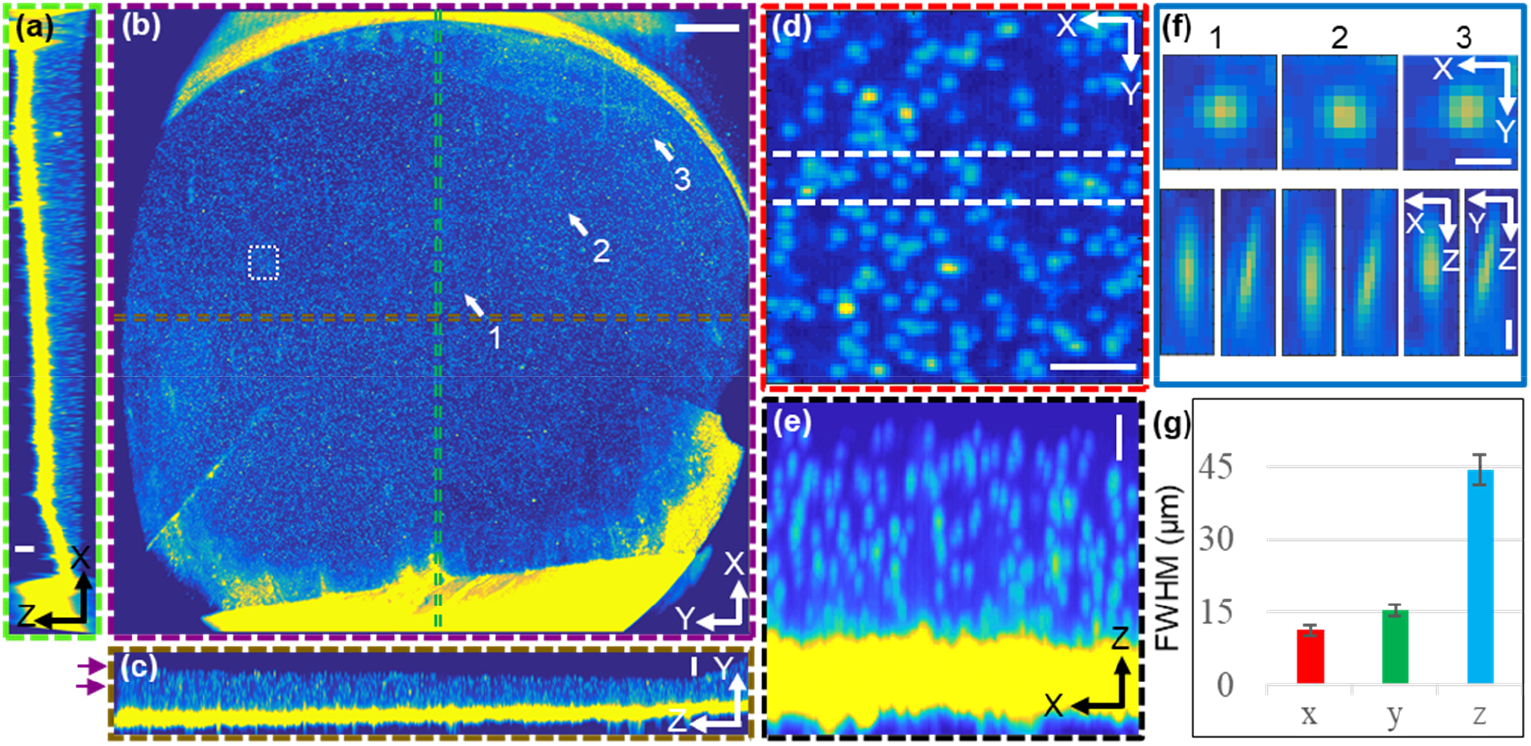
The resolution characterization using a model eye by CoSLO. (a) X-Z cross-section generated by maximum intensity projection (MIP) of the area between two green lines in panel (b); (b) The *en face* view by generated MIP of the area between the purple lines in panel (c) ; (c) The Y-Z cross-section generated by MIP of the area between the brown lines in panel (c); (d) The zoom-in view of the square area in panel (b). (e) The X-Z view of the area between the black dash lines in panel (d). (f) The profile of three representative microspheres. (g) The resolutions along X, Y, and Z directions; Error bar = SD, n=50; Scale bars in (a), (c), (d), (e) are 100 μm, scale bar in (b) is 1000 μm, scale bar in (f) is 25 μm;

**Fig. 4.**
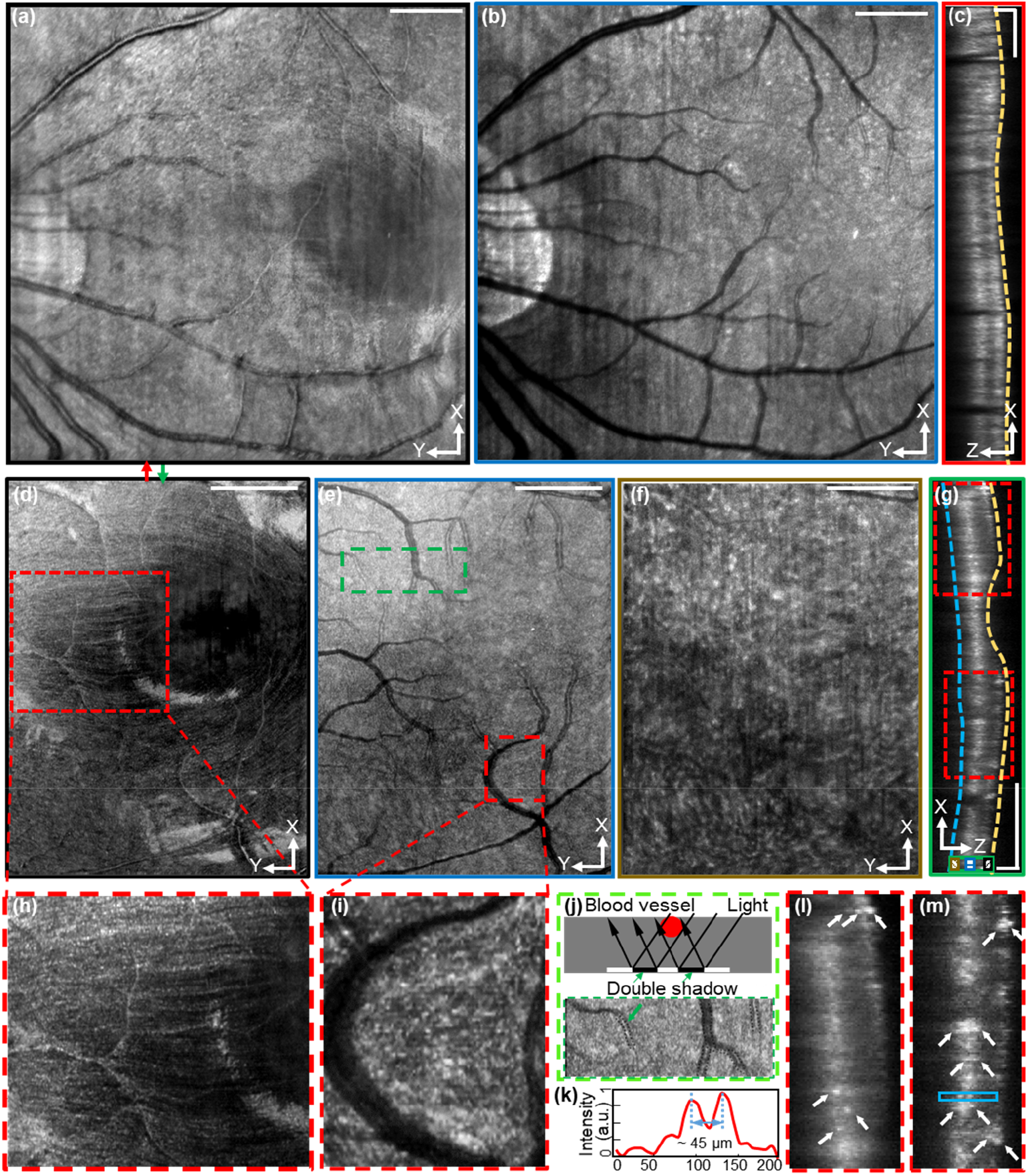
The cross-section images of the human retina by CoSLO. (a-c) and (d-i) are results of subjects 1 and 2, respectively. (a) and (b) are the maximum intensity projection (MIP) of the X-Y planes taken from the superficial layer and the deeper layer indicated by the brown and blue markers in panel (c), respectively. (c) The X-Z cross-section that is taken from the position indicated by the red arrow in panel (a). (d-f) are the MIPs of the X-Y planes taken from the positions indicated by the brown, blue, and black markers in panel (g), respectively. (g) The X-Z cross-section that is taken from the position indicated by the green arrow in panel (d). (h) The zoom-in view of the squared area in panel (d). (i) The zoom-in view of the squared area in panel (e). (j) The explanation on the blood vessels with double edges shown in panel (e). (k)The intensity profile along the depth direction of two adjacent bright spots in the squared area in panel (m). (l) and (m) are the zoom-in views of the squared areas in panel (g). The scale bars for X, Y, and Z direction are 1000 µm, 1000 µm, and 100 µm, respectively.

To observe the details of the retina in 3D, high-definition volume data with a FOV of 2.85 × 4 mm^2^ and a pixel density of 370 × 512 was acquired within 4s. The volume date of subject 2 under the same experiment parameters is shown in supplementary figure S6. Visualization of both volume data can be found in Visualization 2 and 3, respectively. *En face* MIPs from the inner retina, outer retina, and choroid were plotted in Fig. 4(d)-4(f). The depth ranges to generate the *en face* MIPs were indicated in the cross-sectional image in Fig. 4(g). The yellow and blue dash lines indicated the inner and outer boundaries of the retina, serving as depth references. Individual nerve fiber bundles in the superficial retina layer can be clearly observed in Fig. 4(d) and 4(h). The spotty bright signals in Fig. 4(e) and 4(i), which were taken from the outer retina layer, were likely contributed by cone photoreceptors [14]. An interesting observation is the double image of vascular shadow in the *en face* MIPs from outer retina. This is due to the blood attenuations for both the oblique illumination, and the angled detection (Fig. 4j). Features shown in Fig 4(f) were suspected to be signals from the choroid. The three distinct *en face* layers shown in Fig. 3(d-f) is a good indication of the depth sectioning ability of CoSLO.

To better illustrate the depth resolution in CoSLO, we provided Zoom-in views from Fig. 4(g) as in Fig. 4(l) and 4(m). The discrimination of bright contrast is direct evidence of the depth sectioning ability of CoSLO. The normalized A-line intensity in Fig. 4(m) is shown in Fig. 4(k) confirmed that the small feature with a distance of ∼45 µm can be well resolved, which matches previous resolution characterization in the eye model.

In addition to image the central FOV around macula, we tested whether the performance maintains when imaging an off-center FOV on ONH about 6mm away from the fovea in Fig. 5(a)-5(d). The cross-sectional view displayed similar contrast from through the retina as in Fig. 4(c) and 4(g). *En face* projection views were generated within the inner retina, outer retina, and choroid in Fig. 5(b)-5(d). The nerve fiber bundles are clearly visualized within the inner retina where the nerve fiber layer thickens in the peripapillary region (Fig. 5b). The vessel shadow and the rim of the nerve head in Fig. 5(c) are enhanced features in the outer retina (Fig. 4(c)), and we are able to image choroidal vessels in the choroid (Fig. 4(d)) as dark contrast.

**Fig. 5.**
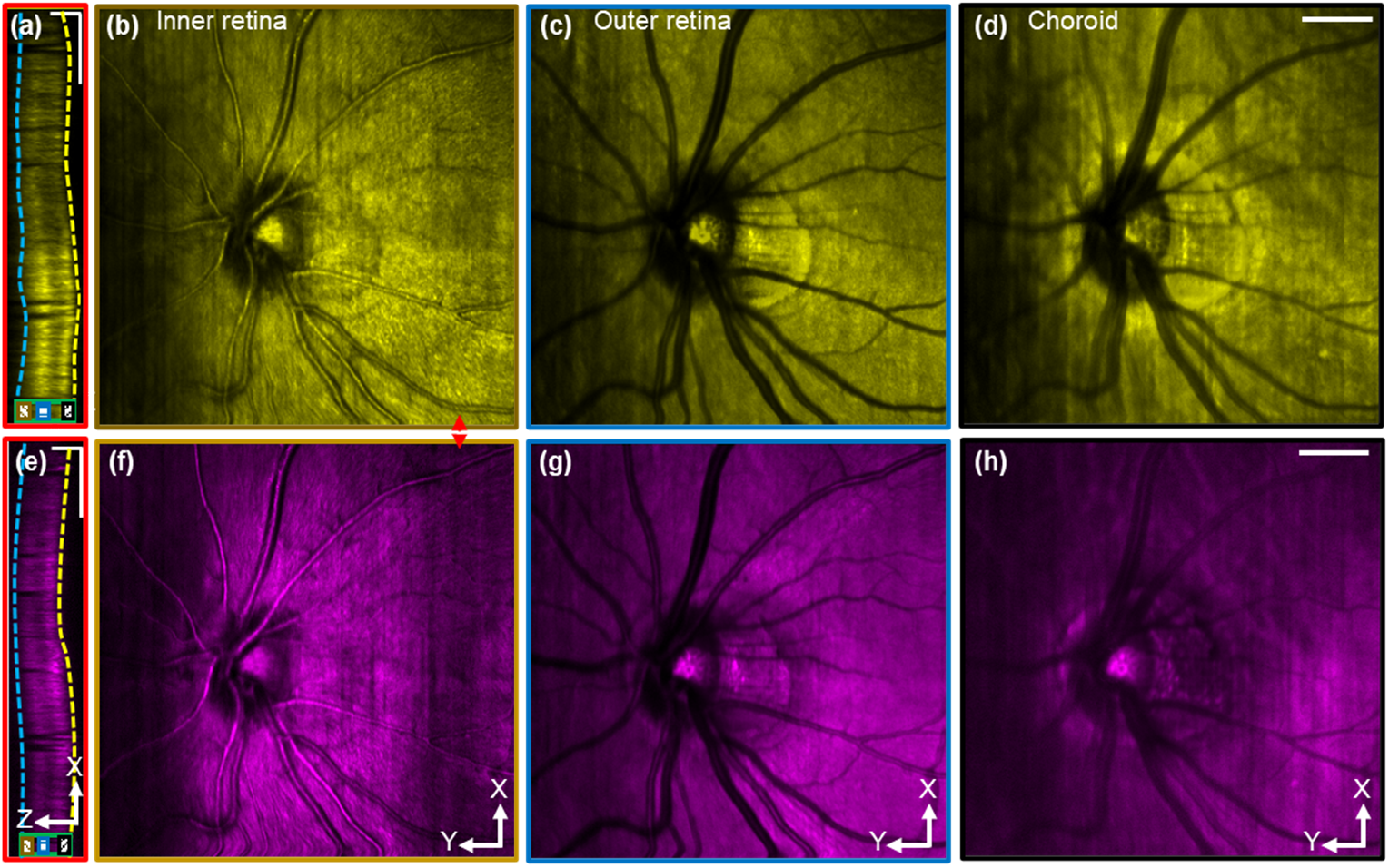
The comparison of the volume data sets acquired by NIR light (a-d) and visible light (e-h). (a) and (e) are the X-Z cross-sectional views taken from the positions indicated by the red arrows in panel (b) and (f), respectively. (b), (c) and (d) are the MIPs of X-Y planes taken from the depth positions indicated by the black, blue, and brown markers in panel (a). The yellow and blue dash lines in panel (a) were the boundaries of the retina that were used as depth references for the segmentation. (f), (g) and (h) are the MIPs of X-Y planes taken from the area indicated by the black, blue, and brown markers in panel (e), respectively. The scale bars for X, Y, and Z direction are 1000 µm, 1000 µm, and 200 µm, respectively.

By simply changing the light source, CoSLO is able to image volumetric spectroscopic contrast in different wavelengths. Figure 5(e)-5(h) shows the CoSLO image with a visible wavelength (600±30nm) illumination on the same OHN FOV. The anatomical features in the cross-section, and three *en face* projections are similar to those with NIR illumination. The signal is relatively weaker in the outer retina and beyond in Fig. 5(e) than Fig. 5(a). The difference could be due to the stronger absorption by photoreceptor pigmentation or RPE in the visible wavelength. Another distinct spectroscopic contrast between NIR and the visible channel is the choroidal vessels, which appear in dark contrast in Fig. 5(d) with NIR illumination, but bright contrast in Fig. 5(h) with visible illumination. Dual-wavelength imaging from a different FOV in subject 2 under the same experiment parameters is shown in supplementary figure S7.)

## 5. Discussion and conclusion

A new 3D imaging modality termed CoSLO for the human retina is presented. It is the first time that large-scale volumetric incoherent imaging is made possible in the living human retina. CoSLO avoids z-stacking and achieves adequate volumetric imaging speed for the human eye by utilizing oblique illumination, remote focusing, and line de-scanning. By utilizing different light sources, wavelength-dependent retina features have been revealed.

We derived the analytical expression for CoSLO, and analyzed how the confocal gating impacts the 3D resolution. At the current setup, numerical simulation suggests 33.5 µm axial resolution, 2.5 and 4.8 µm in X, Y directions, The experimental resolution using microspheres in the gel is characterized to be ∼45 µm axial resolution, 11 and 15 µm in X, and Y. The axial resolution is about four times better than the conventional SLO reported in [12,13] and comparable with AOSLO [14]. The depth resolution in CoSLO is ultimately determined by the accessible spatial frequency range, as shown in the ATF (Fig. 2(a)), particularly in the *k*_*z*_ direction. Although CoSLO only collects the signal existing through the half side of the pupil, the *k*_*z*_ range is largely covered, and therefore the axial resolution is in the order of AOSLO. The experimental resolution is worse than the simulation, presumably due to the aberration. However, since only partial pupil is utilized, CoSLO could be less sensitive to aberration than using the entire pupil, and thus still maintains reasonable axial resolution.

The effect of confocal gating with the linear detector array is similar to that in conventional confocal scanning microscopy, only that the confocal gating is aligned in an angle due to the tilted remote focusing system. Beyond the impact on the diffraction-limited resolution in Fig. 2(g-h), one major benefit of confocal gating is to reject diffusive signal (Supplement 1, Fig. S3) and improve the imaging contrast and resolution. This is particularly useful in imaging highly scattering tissue.

The unique advantage of CoSLO in human retinal imaging is the incoherent volumetric imaging capability that is largely unexplored. A broad range of incoherent contrast can now be harnessed in a 3D manner in the human eye, including exogenous fluorescence contrast (i.e. Fluorescein, indocyanine green) to characterize vascular permeability [32]; intrinsic autofluorescence from photoreceptor [33], RPE [34,35], and choroid [36,37] that has important implications in macular degeneration [38] and other retinal degeneration conditions (e.g. retinitis pigmentosa [39]). Beyond fluorescence, a broad range of incoherent scattering-based contrasts can be explored in 3D in the human retina. For example, the absorption contrast by the photothermal effect to characterize melanin and hemoglobin content [40,41]. The ease of spectral imaging demonstrated in Fig. 5 can facilitate pump-probe methods in the human retina if laser safety is met.

With the successful demonstration of CoSLO in the living human retina, we aim to improve the current system on three major aspects in our ongoing work. First, the imaging speed of our current implementation is mainly limited by the galvanometers which had a large aperture and were used to ensure light collection efficiency. Future improvement can be achieved by using resonance scanner. Second, both the lateral and the axial resolution can be future improved by utilizing adaptive optics. Third, the alignment of the eye pupil will impact the image quality. A motorized chinrest can be used to track and compensate the movement of the pupil in real-time such that the offset of the excitation beam that is projected on the pupil can be well maintained during the acquisition.

In conclusion, CoSLO is a novel method for incoherent volumetric imaging for the human retina. By rapid 3D imaging capability, CoSLO bridges the gap between rich incoherent contrasts and the limitation of the current 2D retinal imaging modalities. We expect CoSLO will be another important imaging method with significant translational values.

## Supporting information

Visualization 1

Visualization 2

Visualization 3

## Acknowledgments

We thank Dr. Zhenpin Guan and Dr. Yahui Wang for their assistance during the experiment.

