## Supplement 1 for "Large field-of-view incoherent volumetric imaging in living human retina by confocal oblique scanning laser ophthalmoscopy"

WENJUN SHAO,[1,2] JI YI[1,2*]

[1]Department of Biomedical Engineering, Johns Hopkins University, Baltimore, Maryland, 21231, USA
[2]Department of Ophthalmology, Johns Hopkins University, Baltimore, Maryland, 21231, USA

### 1. Control signal

The control signal for the galvanometer mirrors and the camera is shown in Fig. S1. GM1 is the fast axis while GM2 and GM3 are operated together as the slow axis. The GM1 was controlled by a sawtooth voltage with 80% duty cycle to create a B-scan in the eye. The camera trigger was synchronized with each step of GM1 in the forward scanning allowing the acquisition of a line image each time. The voltage applied on GM2 is two times that of the GM3 but have a different direction. In this manner, the incident beam projected on GM2 was scanned by both GM2 and GM3 and visually focused on GM1. The voltage of GM2 and GM3 changed at the start of every forward scanning of GM1 such that B-scans at different locations can be acquired.

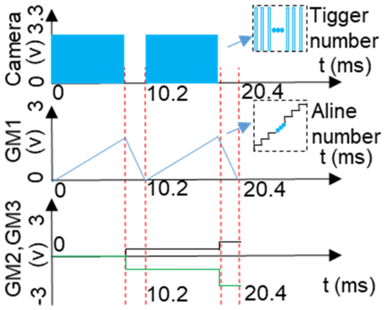


**Fig. S1.** The control signal for the galvanometer mirrors and camera.

### 2. The photograph of the CoSLO

The photograph of the proposed CoSLO is shown in Fig. S2. The setup is equipped with motorized chin rest, fixation target and pupil camera, which will facilitate the fine adjustment and ensure a smooth acquisition process. In addition, the framework of the imaging system is compact, allowing the translation to clinic use with moderate modification.

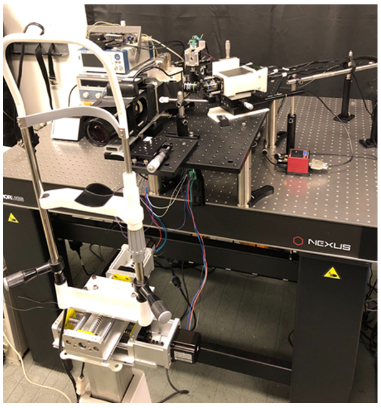

**Fig. S2.** The photograph of CoSLO.

### 3. **The comparison of image contrast under different slit width**

To evaluate the image contrast under different slit widths, the linear detector was replaced by a camera (BFS-U3-51S5M-C, Point Grey) with a two-dimensional CMOS (complementary metal oxide semiconductor) sensor. By binning different numbers of pixels along the column direction of the CMOS sensor, different slit sizes can be simulated. The schematic eye described in [1] model was imaged under different binning configurations. Agarose gel blended with microspheres was adhered inside the schematic eye to mimic the retina. The cross-sectional images of the gel were acquired at the same location under slit widths of 30 µm, 200 µm, and 1400 µm are shown in Fig. S3(a-c). The background of the images in Fig. S3(a-b) is much clear than that of Fig. S3(c). As indicated by the black arrows in Fig. S3(c), it's hard to distinguish the beads from the boundary of the schematic eye due to the diffusive light reflected from the boundary of the schematic eye. However, the beads at the same locations can be separated well from the boundary as shown in Fig. S3(a-b). To quantitatively compare the image contrast, the normalized intensity profiles of three selected beads are analyzed in Fig. S3(d-f). The baseline of the image of the beads acquired under 1400 µm slit width is significantly larger than that acquired under 30 µm and 200 µm. Meanwhile, the image contrast under 30 µm is slightly better than that of 200 µm but very comparable. The full width at half maximum of the bead profiles under 30 µm and 200 slit width µm is comparable which match the simulated results of the MTF very well.

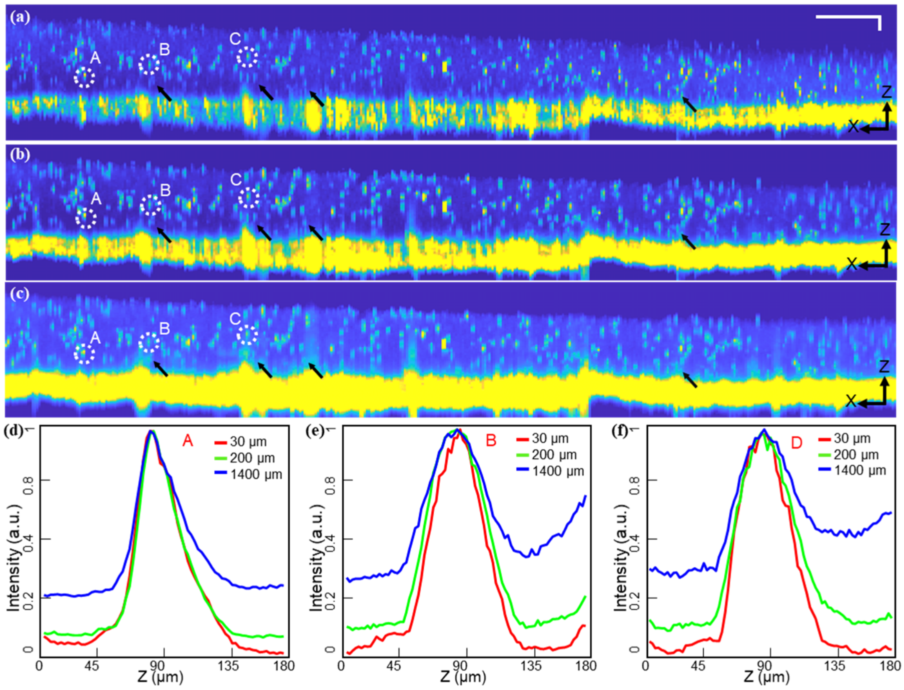


**Fig. S3.** The comparison of image contrast under different slit widths. (a), (b) and (c) are X-Z cross-sectional images of the agarose gel acquired under slit width of 30 µm, 200 µm, and 1400 µm, respectively. (d), (e), and (f) are intensity profiles along the Z-axis of three selected beads that acquired under different slit widths.

### 4. **The calculation of the system parameters**

#### 4.1 The frequency support for the CoSLO

The effective NA of the illumination on the eye can be calculated by $NA_I = 0.5/16$ in which 0.5 mm is the radius of the beam that is projected on the eye, and 16 mm is the equivalent focal length of the eye in the air. Given the ~3 mm offset of the beam that is projected on the eye, the offset of the illumination in the frequency domain is calculated as $k_I = -(2\pi/\lambda)\times 3/16$ . Thus the special frequency range for the illumination is identified as :

$$(k_x)^2 + (k_y)^2 + (k_z)^2 = (2\pi/\lambda)^2, \{(kx, ky) \mid \sqrt{(k_x)^2 + (k_y - k_I)^2} \le (2\pi/\lambda) NA_I\} \tag{1}$$

As the pupil size of the eye is matched with the size of the back aperture of OL1, the equivalent collection NA of the OL2 that mapped on the eye can be calculated as:

$$NA_D = NA_{eye} NA_{obj2} / NA_{obj1} \ , \tag{2}$$

where $NA_{eye}$ is the NA of the eye. The dilated human eye has a NA of~0.2. $NA_{obj1}$ (0.75) and $NA_{obj2}$ (0.55) are the NAs of the OL1 and OL2, respectively. The offset of the detection in the frequency domain can be calculated as:

$$k_D = (2\pi/\lambda)\sin(\theta) NA_{eye} / NA_{obj1} \, , \tag{3}$$

where $\theta$ is the angle between the optical axis of the OL1 and OL2. To make sure the focal plane of OL2 is parallelly aligned with the image in the intermediate image space, $\theta$ was adjusted to be ~ 32°. Given the effective NA ( $NA_D$ ) and the offset frequency ( $k_D$ ) of OL2, the special frequency range for the detection is identified as:

$$(k_x)^2 + (k_y)^2 + (k_z)^2 = (2\pi / \lambda)^2,$$

$$\{(kx, ky) \mid \sqrt{(k_x)^2 + (k_y)^2} \le (2\pi / \lambda) NA_{eye}, \sqrt{(k_x)^2 + (k_y - k_D)^2} \le (2\pi / \lambda) NA_D\} \quad (4)$$

4.2 The size of the virtual confocal aperture

Fig. S4 is a simplified system schematic for the calculation of the size of the virtual confocal aperture. The virtual confocal aperture is the conjugated image of the actual sensor pixel. The width of the actual sensor pixel and the virtual confocal aperture is denoted by $W_{Sensor}$ and $W$, respectively. Given the value of $W_{Sensor}$, the $W$ can be calculated by the system magnification.

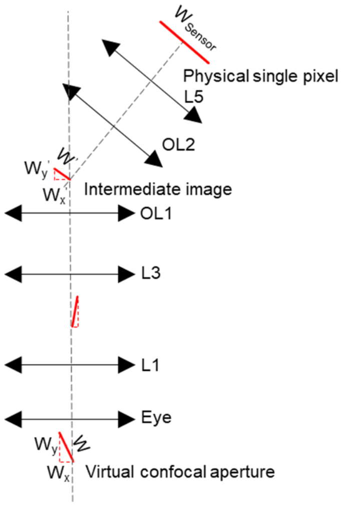


**Fig. S4.** The simplified system schematic for the calculation of the virtual confocal aperture.

As the remote focusing system (OL2, camera, and the linear detector) is angularly aligned, the length of the conjugated image of the actual sensor pixel in the intermediate image space, denoted as $W'$, is calculated firstly. The lateral magnification of remote focusing system can be calculated as:

$$M_{remote} = \frac{f_{L5}}{f_{OL2}}, \quad (5)$$

where $f_{L5}$ (50 mm) and $f_{OL2}$ (3.6 mm) are the focal lengths of the L5 and the OL2, respectively. So, $W'$ is calculated as:

$$W' = \frac{W_{sensor}}{M_{remote}} \quad (6)$$

Secondly, the relationship between the length of the virtual confocal aperture ($W$) and the intermediate image ($W'$) is derived. As the intermediate image of the linear detection is not perpendicular with the optical axis of the OL1, both the lateral and the axial magnification from the virtual confocal aperture to the intermediate image need to be taken into consideration to calculate the size of the virtual confocal aperture. The lateral and the axial magnification from the virtual confocal aperture to the intermediate image, denoted by $M_1$ and $M_2$, can be calculated by their focal lengths as follows:

$$M_1 = \frac{f_{L1} \times f_{OL1}}{f_{eye} \times f_{L3}}, \quad (7)$$

$$M_2 = M_1^{\,2}, \quad (8)$$

where $f_{eye}$ (16 mm) is the focal length of the eye, $f_{L1}$ (50mm) is the focal length of L1, $f_{L3}$ (100 mm) is the focal length of L3, $f_{OL1}$ (9 mm) is the focal length of OL1. Given $M_1$ and $M_2$, the size of the virtual confocal aperture can be calculated as:

$$W_x = \frac{W_x^{'}}{M_1} \tag{9}$$

$$W_y = \frac{W_y^{'}}{M_2} \tag{10}$$

The width vitual confocal aperture can be obtained by:

$$W = \sqrt{W_x^2 + W_y^2} \tag{11}$$

Lastly, by combining Eq. (5-11), the width of the virtual confocal aperture in the sample space can be obtained as $W = W_{Semsor} / 1.3$ . As the height of the virtual confocal aperture is only determined by the lateral magnification, it can be calculated as :

$$H = \frac{H_{Sensor}}{M_{Remote} \times M_1} \tag{12}$$

By combining Eq. (5), (7), and (12), the height of the virtual confocal aperture can be calculated as $H = H_{Semsor} / 3.9$ .

### 5. The imaging of the schematic eye by OCT

To characterize the magnification and the resolution of the proposed CoSLO, a device based on optical coherent tomography (OCT) was used to measure the dimension of the agarose gel adhered at the bottom of the schematic eye [2]. The imaging results are shown in Fig. S5. Fig. S5(a) is the cross-section of the agarose gel. The bright curved edge of the gel indicates the boundary of the schematic eye model. The *enface* view of the gel is shown in Fig. S5(b) in which the circle boundary is the mold that is used to confine the gel from detachment. The radius of the mold was measured to be ~5 mm before the installation in the eye model. The thickness of the agarose gel was measured to be ~0.68 mm. As the OCT device has an axial resolution of ~5 μm, the measured value can be used as ground truth for the resolution calibration of the proposed CoSLO.

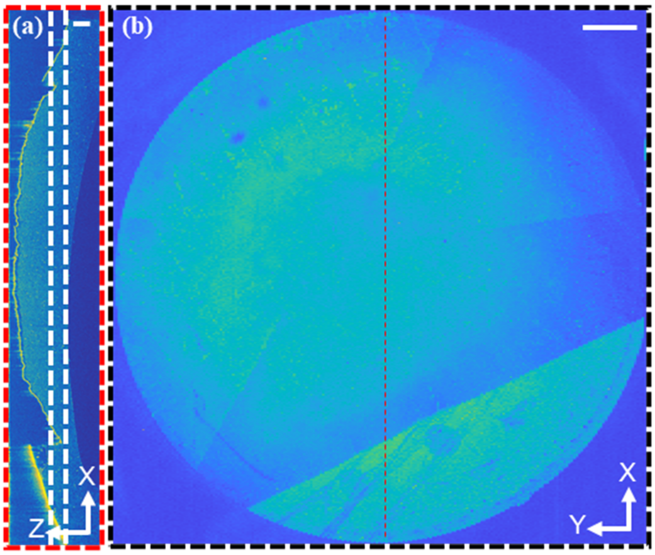


**Fig. S5.** The imaging of the schematic eye by an OCT device. (a) The X-Z cross-section of the agarose gel taken from the position indicated by the red line in panel b. The scale bar is 0.2 mm; (b) The *enface* view of the agarose gel generated by the maximum intensity projection of the layers between the white dash lines in panel (a). The scale bar is 1 mm.

### 6. **Imaging of human subject 2**

The fovea area of human subject 2 was imaged under NIR light. The results are shown in Fig. S6. To generate the *enface* images, the acquired volume data was flatted according to the blue and yellow boundaries shown in Fig. S6(d). The distinctive features in the three different layers shown in Fig. S6(a-c) are a good indication of the depth sectioning capability of the CoSLO. The curvature of the fovea can be observed in the X-Z cross-sectional view in Fig. S6(d), which was taken from the area indicated by the green dash line in Fig. S6(c). The cross-sectional and the *en face* view of the individual nerve fiber bundle can be observed in Fig. S6(e) and (f), respectively. Fig. S6(f) is the zoom-in view of the squared area in Fig. S6(a). The massive bright white spots shown in Fig. S6(g) and (h), which are the zoom-in view of the squared area shown in Fig. S6(b), were suspected to be signals coming from the photoreceptors. Fig. S6(i) is the zoom-in view of the squared area shown in Fig. S6(d). The discrimination of individual spots along the Z-axis is direct evidence of the depth sectioning capability of the CoSLO.

To demonstrate the capability of imaging wavelength-dependent scattering contrast, the fovea area of the human subject 2 was imaged under the NIR light and the visible light separately. The results are shown in Fig. S7(a-c) and Fig. S7(d-f), in which false colors are added to visualize the difference in image contrast. Fig S7(a) and (d) are the X-Z cross-sections taken from the area indicated by the red arrows shown in Fig S7(b) and (e), respectively. The brightness of the outer retina features under these two light sources is different which is probably caused by the different absorption properties of photoreceptor pigmentation or RPE under NIR and visible light. Before the generation of the *enface* view of the inner, and outer retina, the volume data sets acquired under both light sources were flattened according to the blue and yellow boundaries shown in Fig S7(a) and (d). Different textures of the nerve fiber layer around the center of the fovea can be observed when comparing Fig. S7(b) and (e). There are a lot of white spots in Fig. S7(c) while this feature cannot be found in Fig. S7(f). The difference in image contrast could also be a result of different absorption and reflection of the retina tissue under different wavelengths.

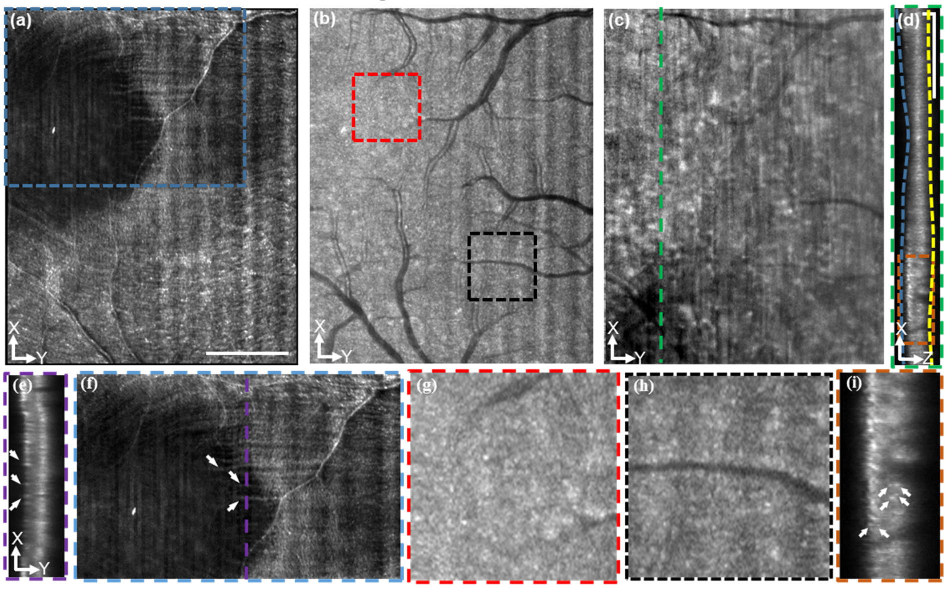


**Fig. S6.** The imaging results of human subject 2 by CoSLO. (a) The maximum intensity projection (MIP) of the ~40 μm along the Z-axis starting from the blue boundary in panel (d). The scale bar is 1 mm; (b) The MIP of the ~50 μm along the Z-axis centered at a position of 110 μm away from the yellow boundary shown in panel (d); (c) The MIP of the ~50 μm along the Z-axis start from the yellow boundary shown in panel (d); (d) The X-Z cross-section that taken from the position indicated by the green line in panel (c).

The vertical and the horizontal scale bars are 1 mm and 0.1 mm, respectively; (e) The X-Z cross-section that is taken from the position indicated by the purple line in panel (f). The cross-sections of the nerve fiber bundles are marked by white arrows; (f) The zoom-in of the squared area in panel (a). Individual nerve fiber bundles are marked by white arrows; (g-h) The zoom-in of the squared area in panel (b); (i) The zoom-in of the squared area in panel (d). Serval white spots in the depth direction are marked by white arrows.

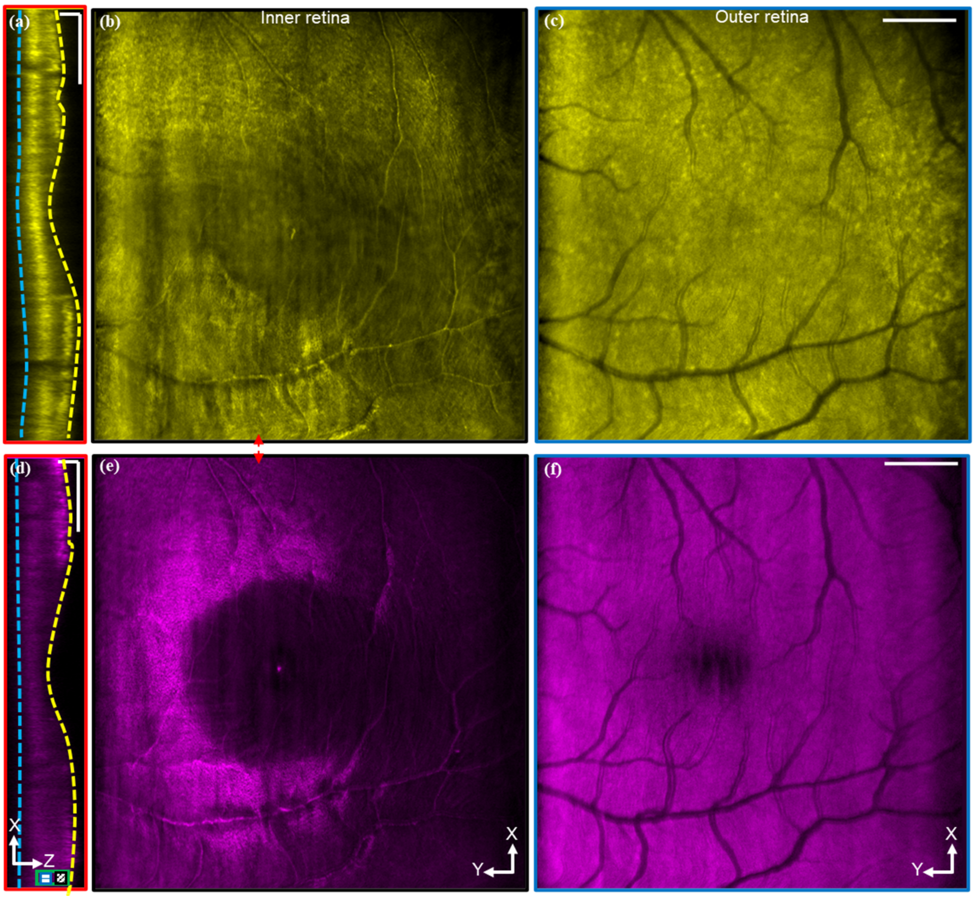


**Fig. S7.** The comparison of the volume data sets acquired by NIR (a-c) and visible light (e-f) on subject 2. (a) and (e) are the X-Z cross-sectional views taken from the positions indicated by the red arrow in panel (b) and (e), respectively. (b) The MIP of ~30 µm along the Z-axis starting from the yellow boundary; (c) The MIP of ~30 µm along the Z-axis centered at a position of 150 µm away from the yellow boundary; (e-f) The *en face* views that extracted by the same method which is used for panel (b) and (c). The vertical scale bar for (a) and (d) is 1 mm. The horizontal scale bar for (a) and (d) is 0.1 mm. The horizontal scale bar for (c) and (f) is 1 mm.
